# Substructure and maturation of lamina-associated domains in neurons of the developing and adult human brain

**DOI:** 10.1101/2024.11.27.625786

**Authors:** Chujing Zhang, Eugene Gil, Sajad Hamid Ahanger, Lynn Mingcong Li, Li Wang, Eric J. Huang, Jingjing Li, Arnold R. Kriegstein, Daniel A. Lim

**Affiliations:** Department of Neurological Surgery; University of California, San Francisco, San Francisco, CA 94143, USA; Department of Neurology; University of California, San Francisco, San Francisco, CA 94143, USA; Eli and Edythe Broad Center of Regeneration Medicine and Stem Cell Research, University of California, San Francisco, San Francisco, CA 94143, USA; San Francisco Veterans Affairs Medical Center, University of California, San Francisco, San Francisco, CA 94143, USA; Department of Pathology, University of California, San Francisco, San Francisco, CA 94143, USA; Department of Pathology and Immunology, Washington University School of Medicine, St. Louis, Missouri

## Abstract

Approximately 30-40% of the human genome is anchored to the nuclear lamina (NL) through variably sized (10 kb-10 Mb) lamina-associated domains (LADs), which can be classified into two subtypes (T1 and T2) based on their level of lamina-association. The dynamics of LAD substructure in cells that remain postmitotic for long periods of time are poorly understood. Here, we developed Genome Organization with CUT and Tag (GO-CaT) to determine the T1- and T2-LAD substructure of postmitotic excitatory neurons isolated from the prenatal and adult human cortex. While T1-LADs exhibited epigenomic features characteristic of stable, cell type-invariant LADs including strong transcriptional repression, in prenatal neurons, T2-LADs were enriched for promoter-enhancer DNA interactions, intermediate levels of gene expression, and genetic risk associated with neurodevelopmental and cognitive disorders. In adult cortical neurons, T1-LADs were expanded in size and genomic coverage, incorporating the majority of the prenatal T2-LADs, sequestering genes involved in neurodevelopment. In contrast, the minority of prenatal T2-LADs that relocated to inter-LAD regions in adult neurons were enriched for processes related to synaptic function. Overall, these data provide evidence that LADs “mature” in postmitotic neurons, remodeling from a genomic architecture that is more permissive for the dynamics of transcription of development to one that is more restricted and focused on the decades-long transcriptional needs of adult brain neurons.

## Introduction

In the interphase nucleus of mammalian cells, chromosomes are radially organized, with specific genomic regions interacting with the lamina of the inner nuclear membrane ^1, 2^. At the nuclear periphery, approximately 30-40% of the genome are segmented into variably sized (10 kb to 10 Mb) lamina-associated domains (LADs), and genes within LADs have low transcriptional activity. In studies of cell differentiation in culture, LADs that “detach” from the lamina exhibit increased gene expression ^3–6^, and conversely, experimental tethering of genes to the lamina suppresses transcription ^7–10^. Thus, the lamina constitutes a large, transcriptionally repressive nuclear subcompartment. Although LADs have been widely studied in cells cultured *in vitro*, relatively little is known about LADs in cells acutely isolated from primary tissues, partly due to limitations of methods used for LAD mapping. In particular, the LAD architecture of post-mitotic cells – such as neurons in the developing and adult brain – is poorly understood.

LADs have substructure based on their degree of lamina-association. Using LaminB1 chromatin immunoprecipitation-sequencing (ChIP-seq) of twelve cultured cell lines, Shah *et al*. classified LADs into two subtypes: T1-LADs, which are highly enriched in LaminB1 and transcriptionally repressed, and T2-LADs, which have moderate levels of LaminB1 enrichment and low levels of gene expression^11^. The potential functional implications of such LAD substructure have not been reported for neurons.

During mammalian brain development, proliferative neural precursor cells give rise to young neurons that at early to mid-gestational ages collect in a region called the cortical plate ^12^. Neurons in the cortical plate are postmitotic but still immature, acquiring additional functions and becoming incorporated into complex circuits later in development. While most cell types are periodically replaced through a process that involves cell division, post-mitotic neurons do not have such an opportunity to globally “reset” their genome architecture through DNA replication and mitosis. Thus, in neurons, the association of chromatin with the lamina may be of particular importance, providing a physical and regulatory scaffold that needs to last a lifetime.

There are several different methods for LAD mapping, each with distinct advantages and disadvantages. Most LAD maps have been generated by DNA adenine methyltransferase identification (Dam-ID) based methods wherein genomic DNA proximal to the lamina undergoes adenine methylation by a transgenic LaminB1-Dam fusion protein and is subsequently identified by sequencing ^13–15^. LaminB1-DamID is not well suited for studying human primary cells due to the need for transgene expression. Other approaches like LaminB1 ChIP-seq ^11^ and Tyramide signal amplification (TSA)-seq ^16, 17^ require tens of millions of cells, limiting their utility for clinical specimens that are usually limited in quantity.

We previously developed Genome Organization with CUT&RUN technology (GO-CaRT), which localizes protein-A micrococcal nuclease (pA-MNase) to LaminB1 antibodies in the nucleus, to map LADs from cells acutely isolated from mouse, macaque, and human brain cells ^18^. In this study, we adapted this technique to utilize pA-Tn5 transposase, providing a streamlined approach with higher sensitivity termed Genome Organization with CUT and Tag (GO-CaT).

Unlike GO-CaRT, GO-CaT uses Tn5-mediated “tagmentation” to insert sequencing adaptors during DNA fragmentation, which greatly improved LAD mapping efficiency and simplified the process. The enhanced LAD mapping efficiency of GO-CaT allowed for precise dissection of LADs into T1 and T2 subdomains from small numbers of neurons isolated from prenatal and adult human brain specimens. Together, these data from human neurons *in vivo* provide novel insights into the substructure of 3D genome architecture and implicate dynamics of T1- and T2-LADs in neuronal differentiation, maturation and human disease.

## Results

### GO-CaT maps LADs with high efficiency

To evaluate the performance of GO-CaT, we first mapped LADs in HEK293T cells, that had been previously mapped by LaminB1 GO-CaRT ^18^ and LaminB1-DamID-seq ^19^. For GO-CaT, we used LaminB1 antibodies to tether pA-Tn5 to the nuclear lamina and identified the tagmented DNA fragments with paired-end sequencing (**Fig.1A**). LADs mapped by GO-CaT were highly similar (Jaccard index 0.68 - 0.84) to those of GO-CaRT and DamID-seq (**Supplementary Fig. 1A, B**). With GO-CaT, under the same sequencing depth, the unique fragments mapped to LADs were 11.3% and 25.1% greater than those of GO-CaRT and DamID-seq, respectively (**Supplementary Fig. 1C**). Furthermore, at LAD boundaries, there was a larger difference of normalized LaminB1 signal within LADs vs. inter-LAD regions as compared to data from GO-CaRT and DamID-seq (**Supplementary Fig. 1D**). These data indicate that the pA-Tn5 activity of GO-CaT efficiently produces accurate LAD maps with discrete LAD boundaries.

**Figure 1.**
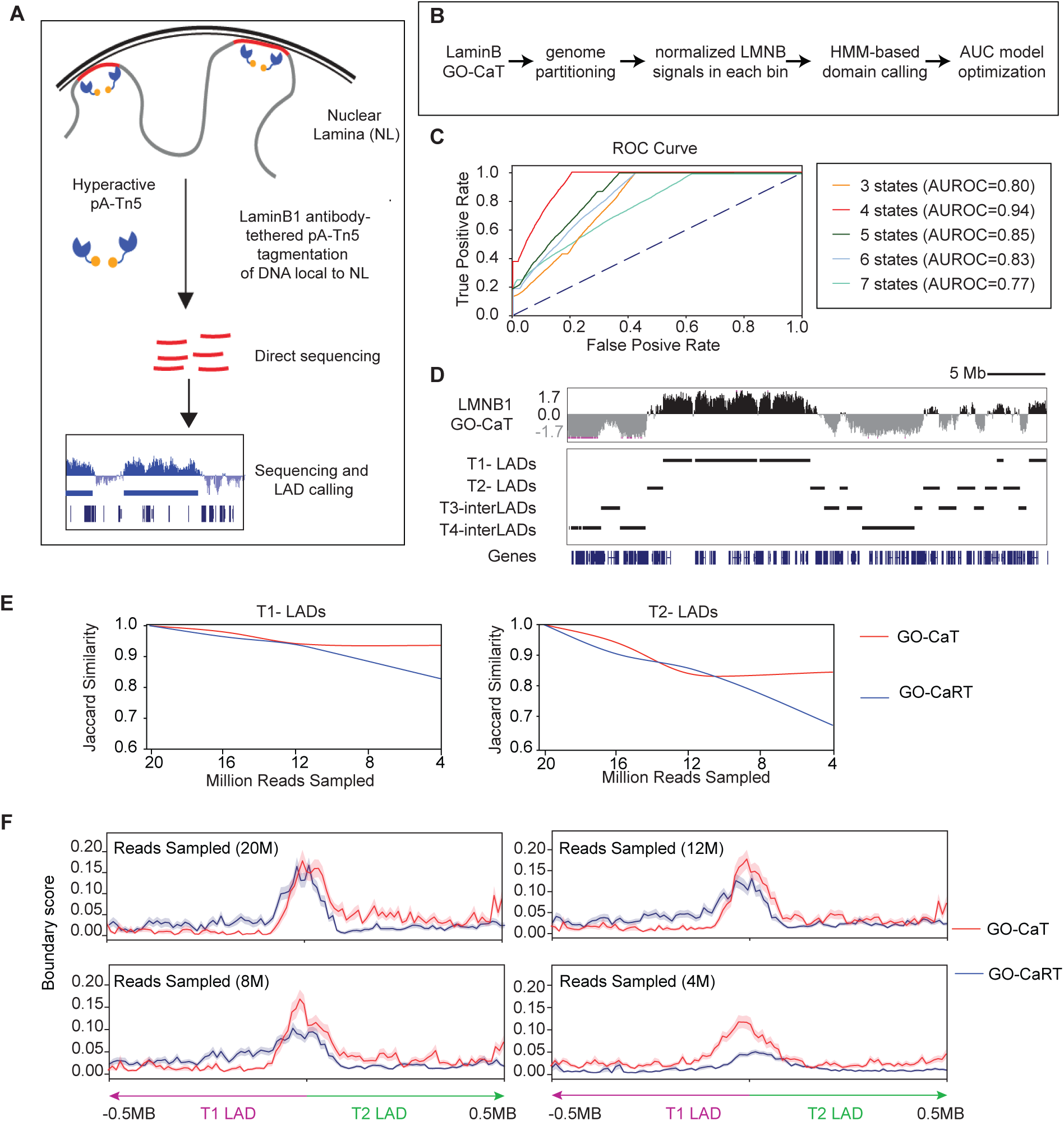
GO-CaT efficiently maps discrete LAD subtypes using a four-state-Hidden Markov Model (HMM) (**A**) Schematic of the GO-CaT method. Lamin B1 antibodies tether hyperactive pA-Tn5 transposase to DNA near the nuclear lamina, enabling targeted tagmentation and sequencing of lamina-associated DNA. (**B**) Workflow for LAD subtypes identification: LaminB1-GO-CaT signal is partitioned genome-wide, normalized per 10Kb bin, and analyzed using an HMM to define chromatin states, optimized using area under the curve (AUC) metrics. (**C**) Receiver operating characteristic (ROC) curves showing HMM performance across models with three to seven chromatin states. (**D**) Genome browser tracks of a representative genomic region (chr7:1 - 41,991,946) illustrating all four identified LAD subtypes in HEK293T cells: T1-LADs, T2-LADs, T3-interLADs, and T4-interLADs, as called by the four-state-HMM using LaminB1-GO-CaT. (**E**) Jaccard similarity plots comparing LAD-subtypes-calling robustness between GO-CaT and GO-CaRT for T1-LADs (left) and T2-LADs (right) when decreasing subsampling rate from 20 million to 4 million reads. (**F**) Metaplots comparing chromatin alternate rate across all T1- and T2-LAD boundaries between GO-CaT and GO-CaRT when subsampling reads from 20 million to 4 million reads. Boundary scores were defined as the averaged transitioning probability from one LAD subtype to another for each 10kb bin predicted by HMM.

### GO-CaT efficiently maps discrete LAD subtypes

Hidden Markov Model (HMM) algorithms have been used to identify LAD subtypes that differ in their levels of LaminB1 enrichment^11^. Using LaminB1 GO-CaT data from HEK293T cells, we partitioned the entire genome into 10kb bins, computed the normalized LaminB1 occupancy for each unit, and used HMM analysis to determine the number of distinct LaminB1 occupancy states that best fit the data (**Fig. 1B**). Receiver Operating Characteristic (ROC) analysis indicated that the four-state model outperformed the three-, five-, six- and seven-state models (**Fig. 1C, Supplementary Fig. 2A-C**). The four-state model segmented the genome into two LADs and two inter-LADs classifications (**Fig. 1D**). Notably, the two LAD states in our analysis corresponded to the previously described T1- and T2-LAD subtypes (**Supplementary Fig. 2D**)^11^. Consistent with prior studies of T1- and T2-LADs across multiple cell types, genes in HEK293T T1-LADs were strongly repressed, whereas genes in T2-LADs displayed moderate expression levels (**Supplementary Fig. 1E**).

To investigate the efficiency of GO-CaT for distinguishing T1- and T2-LAD, we compared GO-CaT and GO-CaRT data, down-sampling the total number of uniquely mapped reads from both datasets to 20, 16, 12, 8 and 4 million and using each subsampled dataset for LAD subtypes identification. With GO-CaT data, LAD subtypes identification was highly consistent across the down-sampled data, having a minimal decrease with fewer reads (**Fig. 1E**). In contrast, with GO-CaRT data, the consistency of T1 and T2-LAD discrimination was decreased as compared to GO-CaT with greater down-sampling (**Fig. 1E**). Additionally, we calculated the genome-wide transition rates at the boundaries of T1- and T2-LADs by the probability gradient computed from the HMM. In GO-CaT data, the transition rates between the two LAD subtypes were very high (transition rate: 0.12) near the boundary even when only 4 million reads were used (**Fig. 1F**). Although the T1- and T2-LAD boundaries were distinctly defined in GO-CaRT when using 20 million reads as an input (transition rate: 0.15), the rate significantly declined after down-sampling, dropping to background levels (transition rate: 0.05) when 4M reads were used (**Fig. 1F**). Taken together, these data highlight the efficiency of GO-CaT in identifying LADs, particularly concerning the distinction of T1- and T2-LAD subtypes.

### GO-CaT identifies LAD subtypes in cells of the human cortical plate

An important advantage of GO-CaT is the highly efficient nature of the technique, enabling the study of relatively small numbers of cells. We therefore employed LaminB1 GO-CaT to identify T1- and T2-LADs of human prenatal cortical development, where only small tissue specimens can usually be obtained. From prenatal brain tissues at gestational weeks (GW) 17 and 22, we dissected the cortical plate regions that are enriched for immature excitatory neurons as previously described^20^. From the cortical plate at GW17 and GW22 we isolated approximately 20,000 cells for LaminB1 GO-CaT (**Fig. 2A**) and utilized the four-state HMM for LAD calling (**Fig. 2B**). The four LaminB1 occupancy states between the two cortical plate samples were highly concordant (**Supplementary Fig. 3A**). Similar to our observations in HEK293T cells, two of these states corresponded to T1- and T2-LADs, exhibiting robust and moderate LaminB1 occupancy, respectively (**Fig. 2B**).

**Figure 2.**
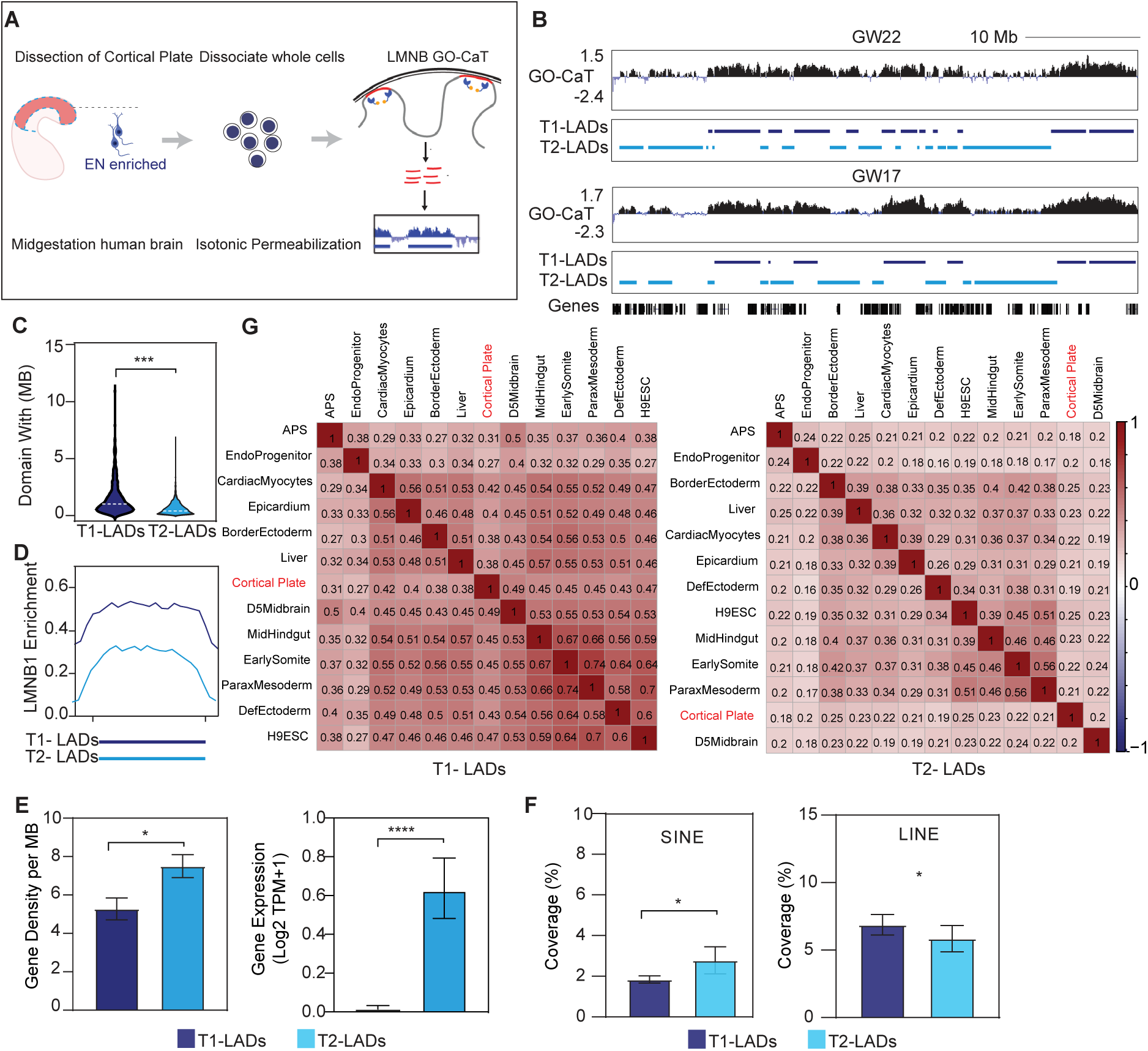
Distinct characteristics of LAD subtypes identified by GO-CaT in the human prenatal brain. (**A**) Schematic of the GO-CaT experimental workflow. Midgestational cortical plate tissues enriched with excitatory neurons were dissected from the human prenatal cortex, followed by single cells dissociation, cell membrane permeabilization, and LaminB1-GO-CaT to map LADs. (**B**) Representative genomic tracks showing T1- and T2-LADs identified by GO-CaT at a shared genomic region (chr1:62,268,975-107,966,486) in GW22 (top) and GW17 (bottom) prenatal brains. (**C**) Violin plot comparing the domain size distribution of T1- and T2-LADs identified in cortical plates (in Mb, million base pairs). (**D**) Metaplot showing the normalized LMNB1 occupancy enrichment over T1-(dark blue) and T2-LADs (light blue). All LADs were scaled to the same size as represented by the solid horizontal bars. (**E**) Bar plots showing (left panel) gene density (number of genes per Mb) and (right panel) average gene expression (log2 TPM) within T1- and T2-LADs as determined by RNA-seq in purified EN. (**F**) Comparison of retrotransposon coverage, showing percentage of regions overlapping with SINEs (left panel) and LINEs (right panel) in T1- and T2-LADs. (**G**) Heatmaps of Jaccard similarity scores for T1-LADs (left panel) and T2-LADs (right panel) across 13 human tissue and cell types. LADs from prenatal cortical plate are highlighted in red, showing greater tissue-invariance of T1-LADs and higher cell-type variability of T2-LADs. *: p < 0.05; **: p < 0.01; ***: p < 0.001; ****: p < 0.0001 (Wilcoxon rank-sum test).

In cortical plate cells, T1- and T2-LADs exhibited similar genome coverage (741Mb and 757Mb, respectively), with each LAD subtype representing ∼24% of the whole genome. Across the genome, T1-LADs were composed of broad domains marked by contiguous LaminB1 occupancy, whereas T2-LADs were characterized by narrower segments interspersed with LaminB1-depleted regions (**Fig. 2B**). Accordingly, the median domain size of T1-LADs (1.2MB) was significantly larger than that of T2-LADs (0.7MB) (**Fig. 2C**). Consistent with previous studies^11^, T1-LADs had higher levels of LaminB1 enrichment as compared to T2-LADs (**Fig. 2D**). T2-LADs of the human prenatal cortex had higher gene density (7.5 vs 5.3 genes per MB) and as expected, greater levels of gene expression as compared to T1-LADs (**Fig. 2E**). Also consistent with prior descriptions of LAD subtypes^11^, T2-LADs in cortical plate cells exhibited greater enrichment of SINE but not LINE transposable elements when compared to T1-LADs (**Fig. 2F**). Together, these data indicate that GO-CaT can accurately map T1- and T2-LAD subtypes in relatively small numbers of cells from human prenatal brain tissues.

We next explored the similarity of LAD subtypes between cultured cells and those of the developing human cortical plate. Comparing an atlas of LADs in differentiated cultured human cell types representing 12 different tissues^11^, we found that T1-LADs of the cortical plate were more similar across different tissues (Jaccard index: 0.27-0.49) as compared to the T2-LADs, which were more dissimilar (Jaccard index: 0.18-0.25) (**Fig. 2G**). Of note, T2-LADs of the developing cortical plate were largely distinct from that of cultured brain cells (Jaccard index: 0.20) (**Fig. 2G**), highlighting the unique LAD architecture of neurons *in vivo* as compared to models of neural differentiation in vitro. Together, these data suggest that T1-LADs are more conserved across different cell types while T2-LADs are more tissue type-specific.

### T1- and T2-LADs of cells in the human cortical plate have distinct epigenomic characteristics

LADs are generally known to consist of heterochromatin enriched with the repressive histone modification H3K9me2, but the epigenomic differences between the T1- and T2-LAD subtypes are less well understood. We performed CUT and Tag analysis for H3K9me2 as well as H3K27me3 in the same cortical plate samples used for GO-CaT LAD mapping. Consistent with studies of LAD subtype in cultured cells^11^, T1-LADs of prenatal cortical plate cells had greater levels of H3K9me2 enrichment as compared to T2-LADs (**Fig. 3A**). Interestingly, T2-LADs were characterized by regions enriched for H3K27me3, which were at lower levels in T1-LADs (**Fig. 3B**). Thus, T1- and T2-LADs have distinct histone modification states in the developing human brain.

**Figure 3.**
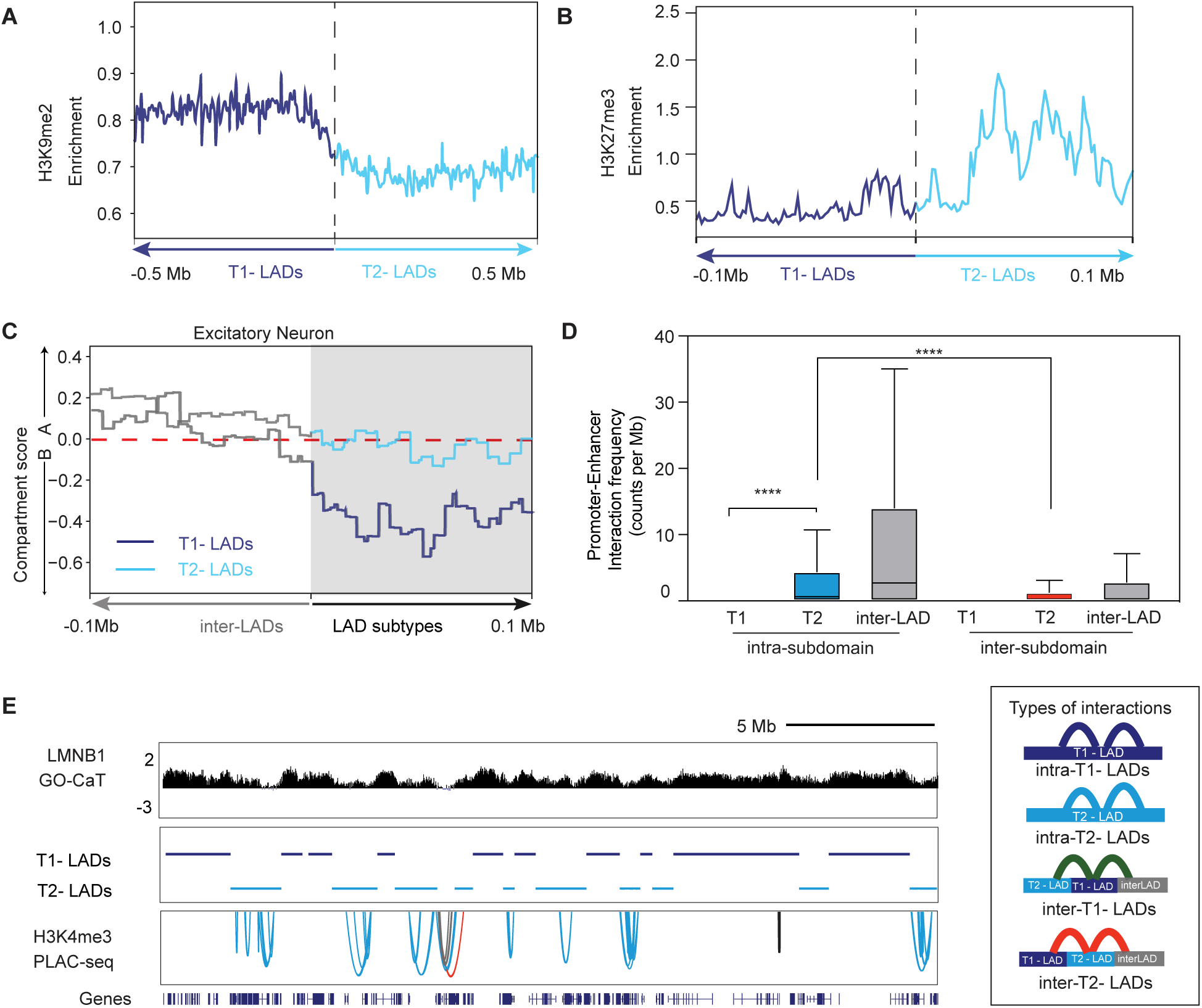
T2-LADs exhibit distinct epigenomic characteristics in the prenatal human cortical plate. (**A, B**) Metaplots showing normalized H3K9me2 (**A**) and H3K27me3 (**B**) enrichment signals across T1-LADs (dark blue) and T2-LADs (light blue) of prenatal cortical plates. All LADs were scaled into a size of 0.5Mb. (**C**) Metaplots showing compartment A/B scores across 100Kb flanking regions of all LAD-inter-LAD boundaries grouped by LAD subtypes (T1: dark blue, T2: light blue). Compartment scores were inferred from the HiC experiment performed in excitatory neurons, and a positive value indicates the state of type A compartment. (**D**) Quantification of promoter–enhancer interaction frequency (counts per Mb) within the same LAD subtypes (intra-subdomain) and crossing the boundaries (inter-subdomain) for T1-, T2- and inter-LADs. T2-LADs exhibit significantly higher intra-subdomain interaction frequency than T1-LADs, though lower than inter-LADs. Adjusted p-values (Wilcoxon rank-sum test with Benjamini– Hochberg correction): ****p < 0.0001. (**E**) Genome browser view of a representative locus (chr4:72,995,600–99,124,210) showing LaminB1 GO-CaT signal, annotated T1- and T2-LADs, and H3K4me3 PLAC-seq interactions. A schematic at right illustrates different intra- and inter-subdomain chromatin interaction types.

DNA-DNA interactions are another important aspect of local chromatin structure^21^, and analysis of DNA-DNA interaction data can broadly separate the genome into two major states called compartment A (more likely to be transcribed) and compartment B (less likely to be transcribed) ^22–24^. We first analyzed high-throughput chromosome conformation capture (Hi-C) data ^25, 26^ from prenatal excitatory neurons in conjunction with our T1- and T2-LAD maps. Across the entire genome, 85.7% and 74.1% T1- and T2-LADs, respectively, were designated as compartment B. However, the magnitude of the compartment scores between T1- and T2-LADs was notably different, with T1-LADs being associated with high compartment B scores, whereas T2-LADs had compartment scores that were intermediate to A and B ^27^(**Fig. 3C**). This observation demonstrates the differential chromosomal conformation of genomic regions across the T1- and T2-LAD subtypes.

To study DNA-DNA looping interactions between promoters and enhancers, we analyzed anti-H3K4me3 PLAC-seq data in excitatory neurons isolated from GW17 to GW22 prenatal brain tissues ^28^. To examine this 3D aspect of local genome organization across the LAD subtypes, we counted the numbers of interactions within each individual T1-, T2- and inter-LAD (*i.e.,* intra-domain DNA-DNA interactions). Notably, we observed a significant enrichment of the intra-domain interactions in T2-LADs as compared to T1-LADs (Fold Change: 10.3, 4.59 per MB vs 0.44 per MB) in the developing cortex (**Fig. 3D, E**). Within the T2-LADs, the intra-domain interactions were more than twice as frequent as long-range interactions across the boundaries of T2-LADs (**Fig. 3D, E**). Thus, the pronounced transcriptional repression of T1-LADs aligns with high levels of H3K9me2, strong association with compartment B, and relative depletion of promoter-enhancer interactions. By contrast, T2-LADs are characterized by lower H3K9me2 levels, representing atypical heterochromatin with higher levels of H3K27me3, intermediate compartment A/B scores and greater levels of promoter-enhancer interactions. Taken together, these data illustrate the unique epigenomic features that distinguish T1- and T2-LADs and emphasize the importance of considering LAD substructures when studying 3D genome organization in developing brains.

### T2-LADs in human cortical plate cells contain genes important for the development of excitatory neurons

The relative enrichment of promoter-enhancer interactions within T2-LADs suggests a transitional state wherein genes possess 3D genomic interactions that may influence their transcriptional activity as cells differentiate. To explore this notion, we first hypothesized that T2-LADs are enriched for genes that are more likely to be neuronal lineage-specific. By analyzing snRNA-seq data in developing human prefrontal cortex ^29^, we mapped gene expression variance across terminally differentiated cell types (upper layer excitatory neuron [EN], deep layer EN, astrocyte and microglia) to T1-, T2-LADs, and inter-LADs identified from cortical plate samples. Genes located in T2-LADs (n=3,986) were nearly twice as variable in terms of expression across these differentiated brain cell types as compared to genes in T1-LADs (n=2,878) and inter-LADs regions (n=29,310) (**Fig. 4A**). These data indicate that genes within the T2-LADs of the prenatal cortical plate are more likely to become expressed in a lineage-type-specific manner as compared to T1-LADs or inter-LAD regions.

**Figure 4.**
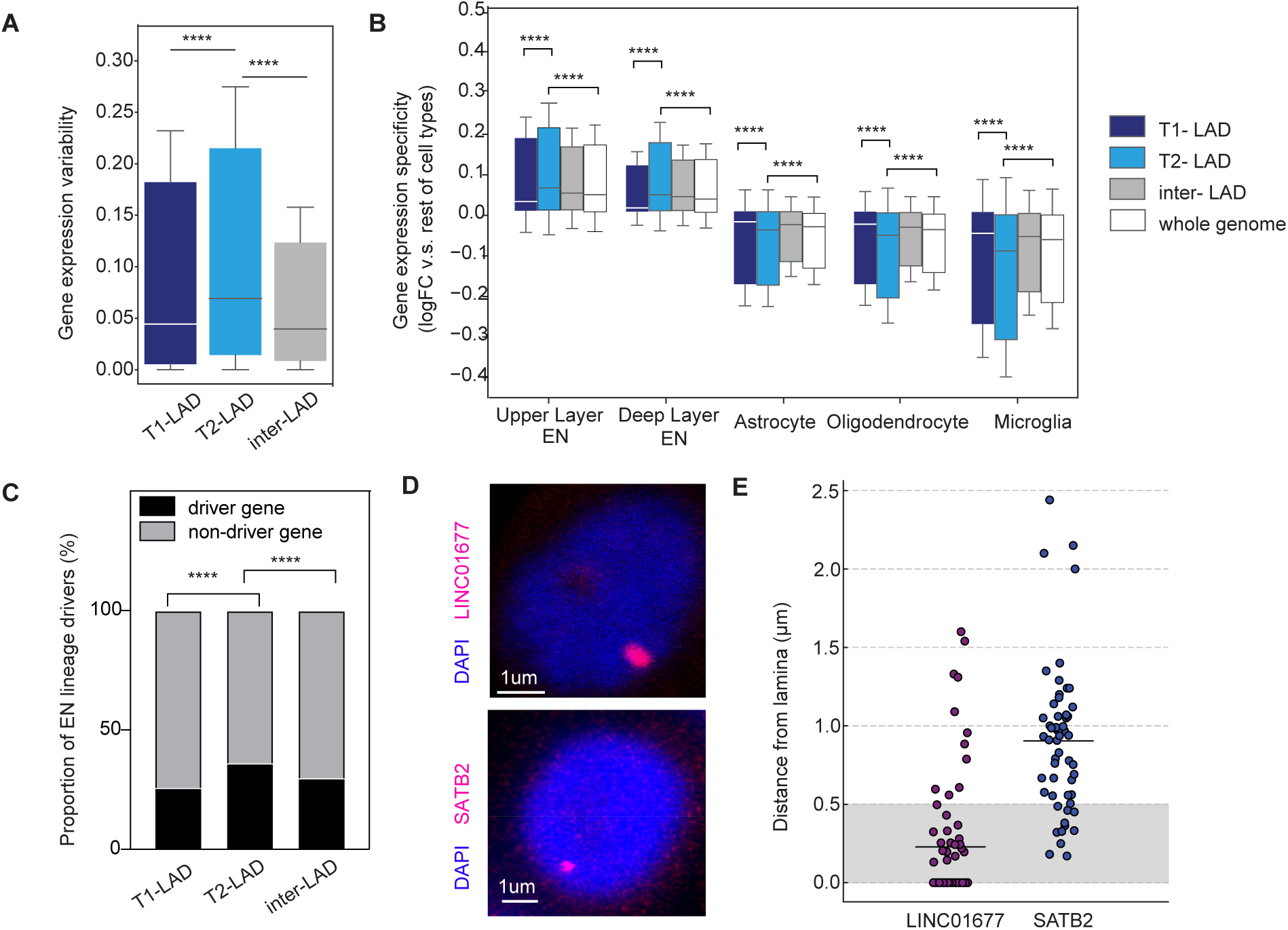
T2-LADs in the human cortical plate contain information important for excitatory neuronal development. (**A**) Box plot showing the standardized expression variability across the genes located within T1-, T2-, and inter-LADs. (**B**) Box plots showing the significantly higher gene expression specificity in EN lineage cell types (Upper Layer and Deep Layer EN) for genes located in T2-LADs compared to those genes from T1-LADs or genome background. Cell type-specific expression was quantified as the log2-fold changes over the mean expression of all other cell types. All P values were from Wilcoxon rank-sum tests and corrected by Benjamini-Hochberg method. **** denotes adjusted P value less than 0.0001. (**C**) Stack barplots demonstrating overrepresentation of neuronal lineage drivers in T2-LADs over T1-LADs and inter-LADs. The proportion of the bar in black denotes the percentage of lineage driver genes inferred from scRNA-seq analysis. All P values were from Fisher’s exact test and corrected by Benjamini-Hochberg method. **** denotes adjusted P value less than 0.0001. (**D**) Representative DNA-FISH images in the nucleus of two representative locus (*LINC01677*, and *SATB2*). (**E**) Box plot quantifying the mean distance of each locus to nuclear lamina.

Notably, genes within T2-LADs of the prenatal cortical plate eventually become expressed and are highly specific to two terminal EN cell types (upper and deep layer) but not to other terminally differentiated states including oligodendrocytes, astrocytes, or microglia (**Fig. 4B**). Because the prenatal cortical plate is enriched for immature ENs ^29^, these observations support the notion that T2-LADs in immature cell types represent a transitional epigenetic state, containing information necessary for subsequent cellular differentiation and function, such as genes that drive cellular lineage specification, identity, and cell type-specific functions. To explore this notion further, we performed pseudotime and lineage analysis using the snRNA-seq data to identify putative lineage drivers of EN differentiation ^30^. We found T2-LADs to be significantly enriched for EN lineage drivers as compared to T1-LADs and inter-LAD regions (**Fig. 4C**). *SATB2* is an EN lineage driver gene with key roles in neuronal development (**Supplementary Fig. 4A**), and DNA fluorescent *in situ* hybridization analysis demonstrated that this T2-LAD gene is located further from the nuclear periphery than T1-LAD gene *LINC01677* (**Fig. 4D, E)**. Taken together, these findings suggest that T2-LADs represent a transitional epigenomic state that contains developmental information important for subsequent neuronal maturation.

### Brain-specific genetic susceptibility variants are enriched in T2-LADs

We next sought to utilize the identified LAD subtypes to interpret genetic variants associated with complex diseases and multigenic phenotypes. We accessed the NHGRI-EBI GWAS catalog ^31^ and selected 176 well-powered GWAS datasets with over 20,000 cases. Many GWAS primarily report only index SNPs with extreme significance, which often fail to capture causal variants as a substantial portion of trait heritability resides in SNPs with associations below the genome-wide significance threshold. To investigate how this heritability is distributed across LAD subtypes, we examined the enrichment of GWAS-associated SNPs within gene loci located in T1-, T2-, and inter-LAD regions via stratified linkage disequilibrium score regression (LDSC) ^32, 33^, a method that quantifies functional enrichment from GWAS summary statistics by leveraging genome-wide SNP data while explicitly accounting for linkage disequilibrium (LD) patterns.

Our analysis revealed a significant difference in per-SNP heritability enrichment across LAD subtypes for multigenic phenotypes and complex diseases (**Table 1**). As expected, risk variants within inter-LADs displayed significant heritability enrichment for 150 complex traits and diseases, but only 15% (26/176) of these traits were neurological phenotypes (**supplementary Fig. 5**). Notably, T2-LAD gene loci demonstrated significant genetic enrichment for phenotypes specific to neuronal development (42.1%, 24/56), whereas no enrichment was observed in T1-LAD loci (**Fig. 5A, B**). For example, risk variants associated with autism spectrum disorder (ASD), putamen volume (Mean Putamen), and attention-deficit/hyperactivity disorder (ADHD) were strongly enriched exclusively within T2-LADs, but not within T1-LADs or inter-LADs (**Fig. 5C**). Additionally, nominal enrichment for non-neurological diseases, such as Type 1 diabetes, was detected only in inter-LADs but not in T2-LADs, supporting the hypothesis that T2-LADs are genetically predisposed to neuronal development disorders and may harbor neuronal-specific regulatory elements.

**Figure 5.**
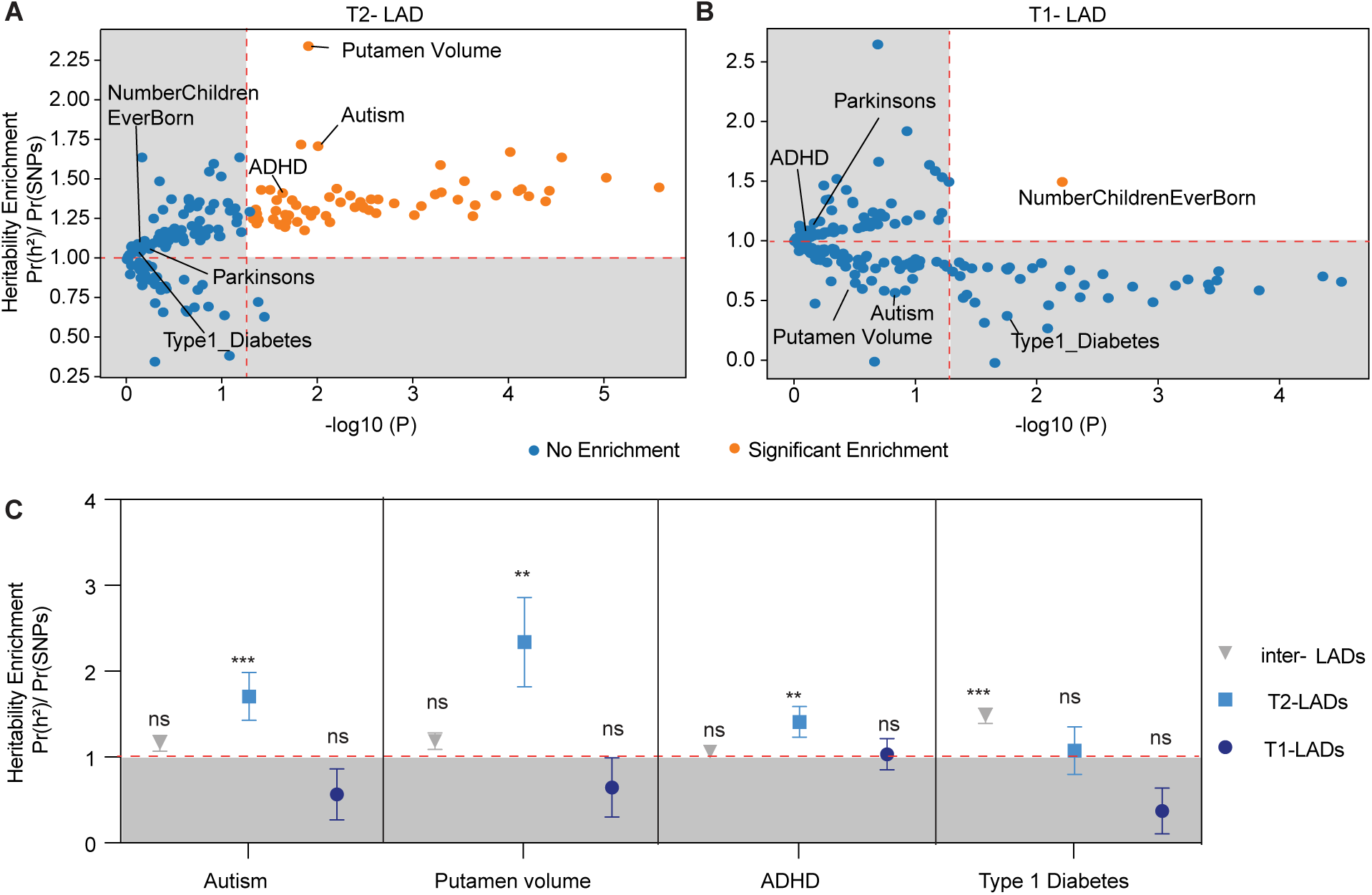
Genetic analysis of LAD subtypes with complex neuropsychiatric disorder-associated variants. (**A, B**) Scatter plots showing the enriched per-SNP heritability from partitioned LDSC analyses for SNPs associated with genes from (**A**) T2-LADs, and (**B**) T1-LADs. X-axis: P values in minus log10 scale; Y-axis: Enrichment of heritability for each SNP. Each dot represents a GWAS phenotype tested and is color-coded by their significance of enrichment. (**C**) Comparisons of per-SNP heritability enrichment between genetic variants associated with inter-LADs, T2-LADs or T1-LADs on selected complex traits and multigenic diseases (e.g. autism, mean volume of putamen, ADHD, and type I diabetes). ***: P values < 0.001; **: P value < 0.01; ns: not significant.

### Extensive remodeling of LAD subtypes occurs during neuronal maturation

Given that T2-LADs in prenatal neurons appeared to represent a transitional epigenomic state related to neuronal development, we hypothesized that the LAD architecture of adult brain neurons would be substantially different, expanded with T1-LADs and reduced with T2-LADs. Although the cortical plate of the prenatal cortex is composed of mostly ENs, there are also a smaller number of inhibitory neurons (INs). To reduce potential heterogeneity of the cell population analyzed, we purified excitatory neurons (EN) from prenatal cortical plate tissues of three individuals (GW18, GW20 and GW22) by fluorescent activated nuclei sorting (FANS) for SATB2-expression and performed LaminB1-GO-CaT (**Fig. 6A**, **Table 2**). The LAD substructures identified from purified prenatal ENs recapitulated those of the developing cortical plate (**supplementary Fig. 6A**).

**Figure 6.**
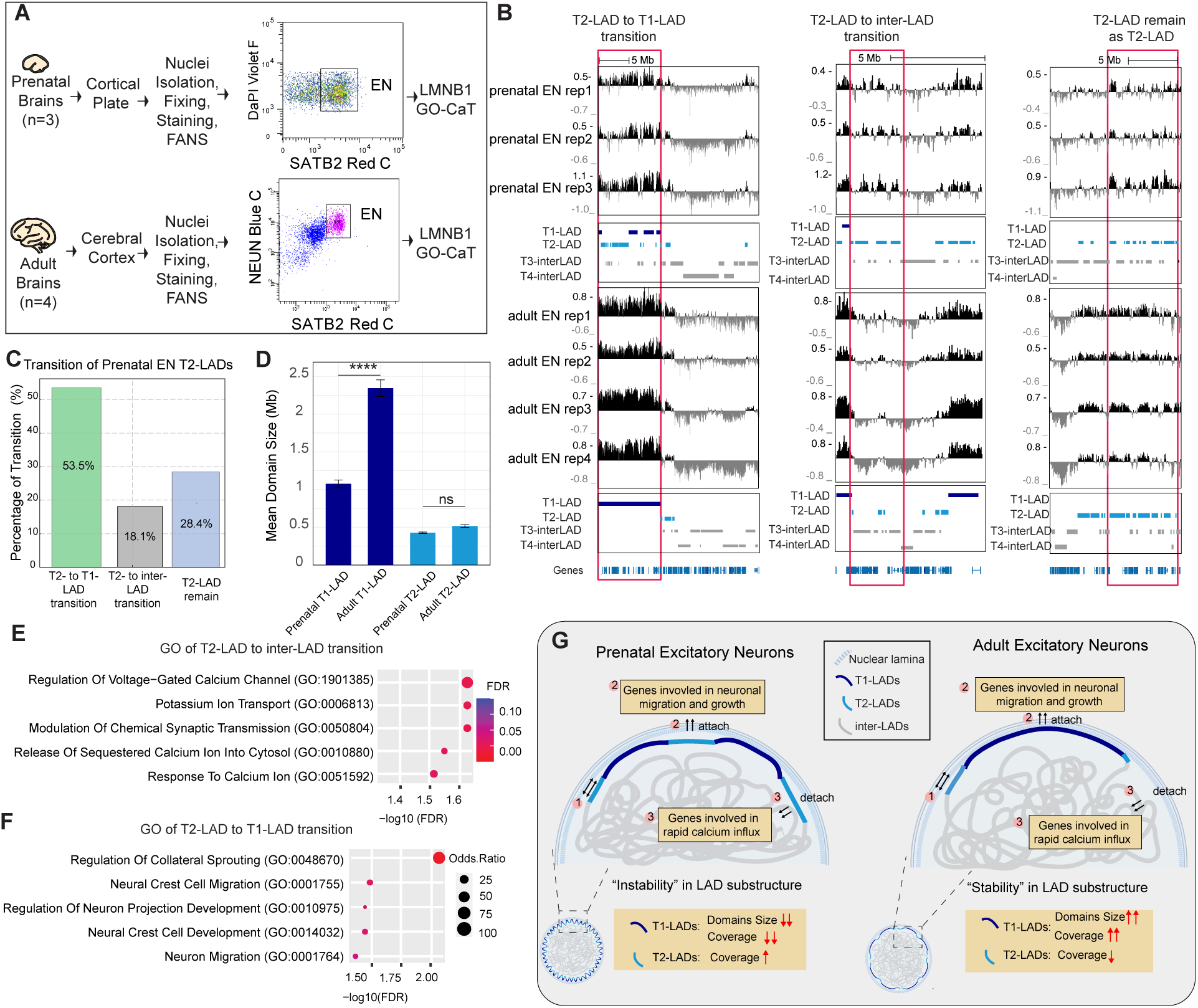
LAD subtypes are extensively remodeled during human neuronal maturation. (**A**) Schematic of the sorting strategy of prenatal and adult EN. (**B**) Genomic tracks illustrating transitions of LAD subtypes during EN maturation. The genomic tracks show four distinct LAD subtypes (T1-LAD, T2-LAD, T3-inter-LAD, and T4-inter-LAD) identified using GO-CaT across representative genomic regions. These regions exemplify the transition of T2-LADs to T1-LADs (left panel: chr1:213,396,947–240,027,663), T2-LADs to inter-LADs (middle panel: chr18:45,203,729–53,922,055), and stable T2-LADs that remain unchanged during maturation (right panel: chr5:149,184,675–163,357,681). Scale bar represents 5 Mb. (**C**) Bar plot showing the percentage of prenatal EN T2-LADs that either transition to T1-LAD, inter-LAD or remain stable as T2-LAD during neuronal maturation. Percentages of each transition state are indicated within the corresponding bars. (**D**) Mean domain size of T1- and T2-LADs in prenatal and adult excitatory neurons (EN). Error bars represent standard error of the mean (SEM); **** p < 0.0001; ns, not significant. (E, F) Dot plots of gene ontology analysis for genes transitioning from T2-LADs in prenatal EN to T1-LADs in adult EN (E), from T2-LADs in prenatal EN to inter-LADs in adult EN (F). FDR (adjusted P values) are color-coded, and size of the dots represents the enrichment odds ratio. (**G**) Graphic summary of LAD substructure remodeling and stabilization during human neuronal maturation. In adult EN, T1-LADs expand in size and genomic coverage, largely replacing T2-LADs that were previously embedded between T1-LADs in prenatal neurons, while overall T2-LAD coverage decreases concurrently. This shift strengthens genome-nuclear lamina attachment and supports a less disrupted and more stable nuclear periphery, as visualized in the zoomed-out views of the nuclei (left corner panels). The remodeling of LAD substructure is closely associated with biological functions during neuronal maturation: genes involved in neuronal migration and growth relocate from T2- to T1-LADs for transcriptional repression, while genes involved in calcium signaling which reflects the increased functional demands of adult neurons for rapid calcium influx to support synaptic transmission and complex processes like learning and memory, relocated from T2-LADs to inter-LAD regions during neuronal maturation.

To investigate LAD architecture in mature human neurons, we performed FANS to purify SATB2+ EN from postmortem adult cerebral cortex tissues (middle frontal gyrus and superior temporal gyrus) collected from four individuals (**Fig. 6A**, **Table 2**). We applied LaminB1-GO-CaT to the isolated nuclei, which also expressed the neuronal marker NeuN, and used our four-state HMM to map both T1- and T2-LAD subtypes in adult EN (**Fig. 6A, B)**.

Strikingly, the genomic coverage of T1-LADs in adult EN (933 Mb, 33.2% of the whole genome) was more than twice that of prenatal neurons (416 Mb, 14.8% of the whole genome). This expansion of T1-LADs was primarily driven by a large-scale reorganization in which 53.5% of T2-LADs in prenatal EN (407 Mb, 14.5% of the genome) were relocated into T1-LADs in the adult EN (**Fig. 6C**). These transitioning prenatal T2-LADs were frequently bordered on both sides by T1-LADs in prenatal EN (**Fig. 6B**), resulting in a ∼2.2-fold increase in the average domain size of T1-LADs in adult EN (**Fig. 6D).**

A smaller proportion of prenatal T2-LADs (137 Mb, 4.9% of the genome) relocated to inter-LAD genomic regions in the adult EN, further contributing to the reduction in T2-LAD genomic coverage during neuronal maturation (**Fig. 6B, C**). In contrast to the highly dynamic nature of prenatal EN T2-LADs, the majority of T1-LADs remained stable between prenatal and adult neuron populations: ∼64.8% of prenatal T1-LADs (270 Mb, 10.0% of the genome) were retained in neurons of the adult cerebral cortex (**supplementary Fig. 6B**). Only a small proportion of prenatal inter-LAD regions were relocated to T1- and T2-LADs in adult EN (∼16.1% and ∼18.0%, respectively) (**supplementary Fig. 6B**). Together, these transitions reflect a large-scale reorganization of LAD substructure, characterized by a shift toward an expansion of T1-LADs and a reduction of T2-LADs. This remodeling results in a more consolidated lamina-genome association in adult EN, suggesting enhanced spatial genome stability at the nuclear periphery (**Fig. 6G**).

To characterize the expression level of genes that relocated from T2-LADs to T1-LADs and inter-LAD regions, we analyzed snRNA-seq data from Herring et al., which profiled excitatory neurons across 24 developmental time-points from GW22 to 40 years of age ^29^. To quantify the rate of gene expression change over development, we fit expression trends for each gene to a regularized regression model. Genes that transited from T2- to T1-LADs in adult EN exhibited lower levels of gene expression (**supplementary Fig. 6C**). In contrast, genes that relocated from T2-LAD to inter-LAD regions exhibited significantly increased expression levels compared to those that remained within T2-LADs (p=5.92e-7) or those situated in regions transitioning from T2-LADs to T1-LADs (T2-to-T1 genes, p=1.27e-6) (**supplementary Fig. 6C**). Notably, genes that transitioned from T2-LADs to inter-LADs showed a gradual increase in expression over time during neuronal maturation, whereas relocation to T1-LADs was associated with transcriptional repression (**supplementary Fig. 6D**). This dynamic expression pattern was not observed in snRNA-seq data from IN or oligodendrocytes (**supplementary Fig. 6C**), supporting that the observed chromatin architecture and gene expression changes are specific to ENs in the adult brain.

Genes that relocated from prenatal EN T2-LADs to inter-LAD regions in adult EN (e.g., *CHRM3* and *DMD*) were enriched in gene ontology (GO) terms related to calcium signaling responses (GO: 1901385, FDR=0.024; GO: 0051592, FDR=0.031) (**Fig. 6E, supplementary Fig. 6E**), supporting previous findings of increased functional demands of human neuronal maturation for rapid calcium influx to support synaptic transmission and complex processes like learning and memory ^34–37^. While T1-LADs found in both prenatal and adult brain were not enriched for any GO terms, genes that relocated from T2- to T1-LADs in adult EN (e.g., *SEMA6D* and *LPAR1*) were enriched for GO terms related to neuronal migration and growth (GO:0014032, FDR=0.036; GO: 0001764, FDR=0.045) (**Fig. 6F, supplementary Fig. 6F**), which are essential for early postmitotic neurons but are less involved after neural circuits are established ^37–40^. Together, these findings suggest that LAD substructure matures across human neurodevelopment, underlying a shift away from developmental functions to those that are specialized to the functions and demands of fully differentiated neurons (**Fig. 6G**).

## DISCUSSION

Although LADs have been studied in numerous different cell lines in culture ^11^, there are very few reports of this aspect of 3D genome architecture in cells *in vivo*. Leveraging the advantages of Tn5 and related CUT&Tag methods, LaminB1 GO-CaT exhibited greater efficiency for LAD mapping, enabling the study of relatively small numbers of cells acutely isolated from tissues. Recently, pA-DamID has been introduced for LAD-mapping and likely provides similar advantages^14^. Using GO-CaT, we identified the LAD substructure of neuronal cell populations from the developing and adult human brain. Our studies of T1- and T2-LADs in the mid-gestational cortical plate revealed novel insights into the potential roles of LAD substructure in neurogenesis and phenotypes related to genetic variation. Mapping LAD substructure in a population of adult brain neurons provides foundational groundwork for understanding how spatial 3D genome architecture can mature in cells that remain postmitotic for decades.

For LAD mapping, LaminB1 GO-CaT has performance characteristics that make it well-suited for experimental scenarios in which samples are limited (*e.g.* clinical specimens and FACS-sorting of rare cell populations). In our benchmarking studies, GO-CaT mapped LADs with high confidence from as few as 10,000 cells. Subsampling analyses revealed LaminB1 GO-CaT to have the lowest marginal input cell requirement as compared to DamID- and GO-CaRT-based methods. Furthermore, with GO-CaT data, T1- and T2-LAD subdomains had clearer boundaries, resulting in higher resolution and lower background during LAD calling.

In addition to confirming previously described key characteristics of LAD subdomains, we discovered additional epigenomic characteristics that distinguish T1- and T2-LADs in prenatal cortical plate cells. Consistent with prior observations made in cultured human cell lines ^11^, T1-LADs in primary neuronal cells exhibited high levels of H3K9me2 and correspondingly low gene expression. Adding to the understanding of T2-LADs being a less repressed LAD state, we found that T2-LADs in prenatal brain cells have relatively higher levels of H3K27me3, greater association with compartment A, and more frequent DNA interactions between promoters and potential distal regulatory sequences. Collectively, this epigenomic signature of T2-LADs suggests that this LAD substructure represents a distinct chromatin state that may be poised for transcriptional activation.

The higher levels of DNA-DNA interactions with promoters in T2-LADs suggests a chromatin configuration that allows genes to be activated in a lineage-specific manner as they differentiate. Genes located in T2-LADs demonstrated higher variability in expression across differentiated brain cell types as compared to T1-LADs and inter-LADs regions, indicating that T2-LADs harbor lineage-specific genes essential for neural cell identity. These genes were enriched for neuronal-specific activities, such as neuronal projection development (GO:0010976, FDR=0.0014; GO:0048666, FDR=0.0031), while T1-LADs genes were related to more general cellular functions like cell-cell adhesion and cell cycle (GO:0098742, FDR=9.39E-04; GO:0090068, FDR=0.0049). Overall, genes in T2-LADs were predominantly activated in terminally differentiated ENs, emphasizing the role of T2-LADs in encoding information relevant for EN lineage specification.

Our integrative genome analyses revealed prenatal T2-LADs exhibit higher genetic susceptibility of genome variations related to neurodevelopmental disorders like ASD and ADHD. In contrast, inter-LADs were more broadly associated with diverse human phenotypes but had limited relevance to neurological traits, further emphasizing the specificity of T2-LADs to brain-related genetic predispositions. These findings suggest that the unique epigenomic state of T2-LADs represents a key interface between genetic risk and cellular function in developing neurons.

In adult neurons, the landscape of LADs was substantially reorganized, as compared to prenatal cells. Most notably, adult neurons have a reduction in T2-LADs. This shift appears to reflect a maturation process wherein T2-LADs are repositioned to more distinct chromatin states, either transitioning to inter-LAD regions or T1-LADs. Genes that relocated from T2- to T1-LADs in adult neurons exhibited reduced gene expression, consistent with the role of T1-LADs as repressive chromatin domains that help maintain a transcriptionally restrained environment. The observed 27.1% increase in T1-LAD coverage may signify the establishment of a more differentiated genome architecture, supporting neuronal function while restricting the processes that are more relevant to earlier stages of neuronal maturation. The relocation of T2-LADs to inter-LAD regions in adult neurons corresponded to increased gene expression, suggesting a transition from a poised to active epigenomic state of genes related to adult neuronal function. More broadly, the comparison of prenatal and adult neuron LAD substructure provides insights into how the spatial genome architecture may “mature” in postmitotic cells to achieve a long-lived, cell-type specific transcriptional regulatory scaffold.

**Supplementary. Figure 1.**
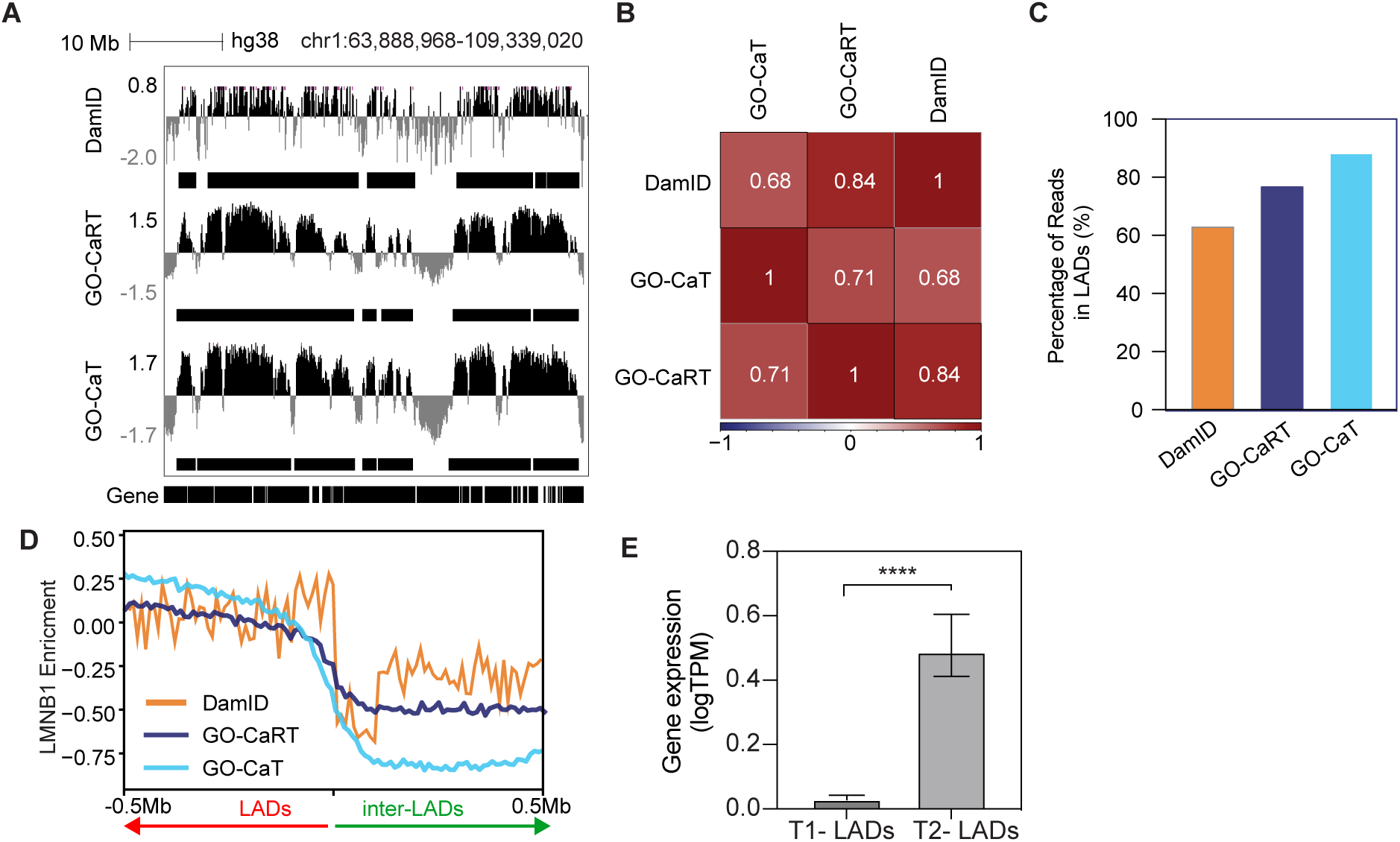
GO-CaT is an efficient method for profiling genome-wide LAD structure. (**A**) Genome tracks comparing LaminB interaction profiles captured by LaminB DamID, GO-CaRT and GO-CaT in HEK293T cells at a representative region (chr1:63,888,968-109,339,020). The Y axis represents the log ratio of LaminB signals over negative controls for each experiment. Identified LADs were denoted by solid horizontal bars. (**B**) Correlation heatmap showing the Jaccard similarity between LADs identified by LaminB1 DamID, GO-CaRT and GO-CaT in HEK293T cells. (**C**) Percentage of sequenced reads in LADs identified by DamID (orange), GO-CaRT (dark blue) and GO-CaT (light blue) in HEK293T. All experiments were sub-sampled to have equivalent number of reads and sequencing depth for comparison. (**D**) Metaplot showing normalized LaminB1 enrichment across all 0.5Mb flanking regions of the boundaries that distinguish LADs from inter-LAD in HEK293T. (**E**) Bar plot showing genome-wide gene expression difference between T1- and T2-LADs in HEK293T cells. P value was determined by Wilcoxon rank sum test, and **** denotes a p value less than 0.0001.

**Supplementary. Figure 2.**
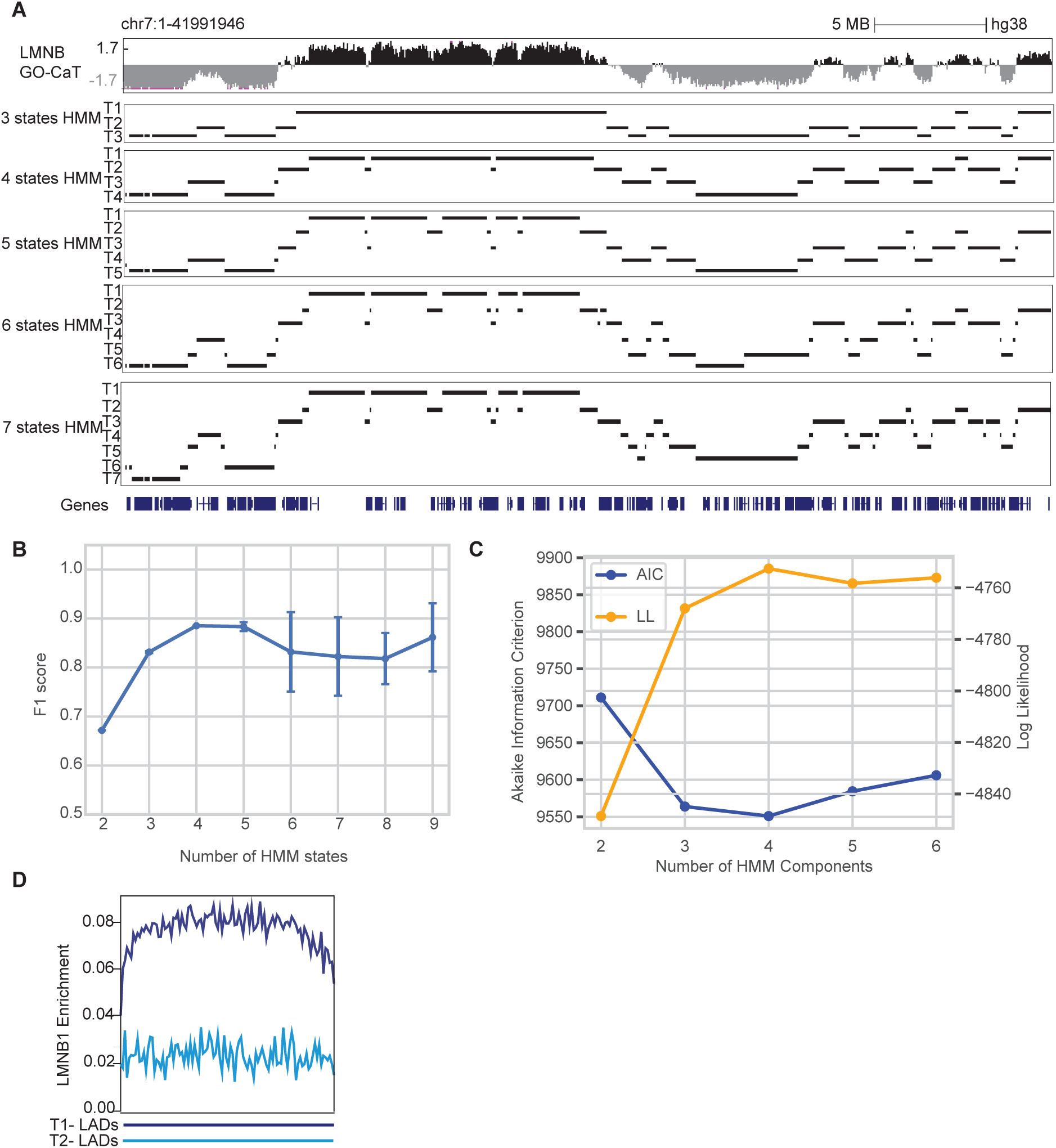
Comparison of HMM performance of different states of HMM on mapping LADs subtypes. (**A)** Genome tracks showing the different numbers of partitioned chromatin states (n = 3 to 7) identified by HMM at a representative genomic locus in HEK293T cells. (**B)** Performance evaluation by F1 score with given numbers of states used in HMM. F1 scores were derived from precision and recall when benchmarking the called LADs with the LB1 occupancy enrichment across all 10kb bins. (**C)** HMM model selection with optimal number of states by AIC (Akaike Information Criterion) and log-likelihood (LL). The lowest AIC and the highest LL indicate the best model fit. (**D)** Normalized LMNB level over T1-(dark blue) and T2-LADs (light blue) in HEK293T cells scaled to the same relative size depicted by a solid horizontal bar.

**Supplementary Figure 3.**
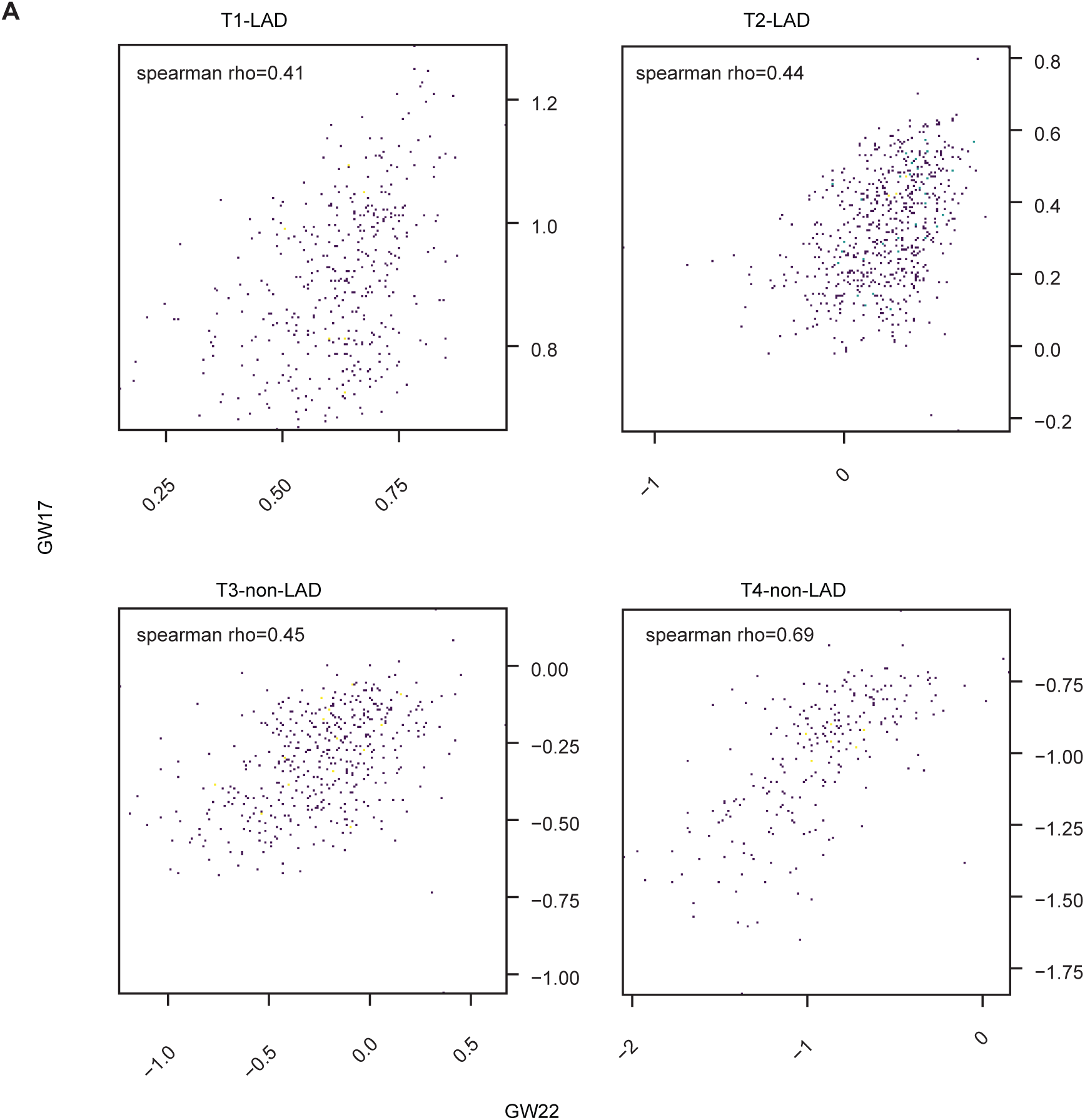
LAD subtypes are highly replicable in different primary tissues. Scatterplots showing the Spearman correlation of LB enrichment over four subtypes of chromatin states between biological replicates of primary cortical plates from two donors of different gestational ages (GW17 v.s. GW 22).

**Supplementary Figure 4.**
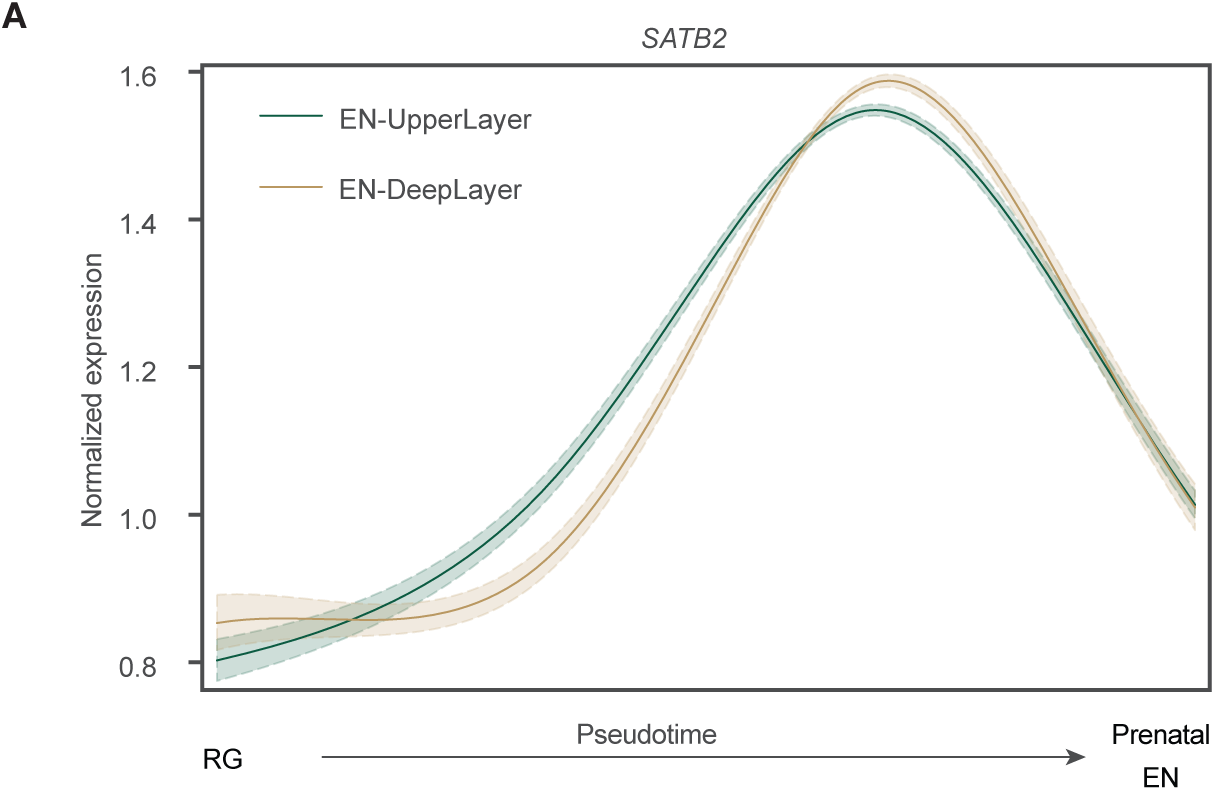
Expression of SATB2 over developmental time. The line plot represents the smoothed expression trend of SATB2 expression for each cell ordered by the reconstructed developmental trajectory from RG (radial glia) to EN (excitatory neurons) captured in snRNA-seq data.

**Supplementary Figure 5.**
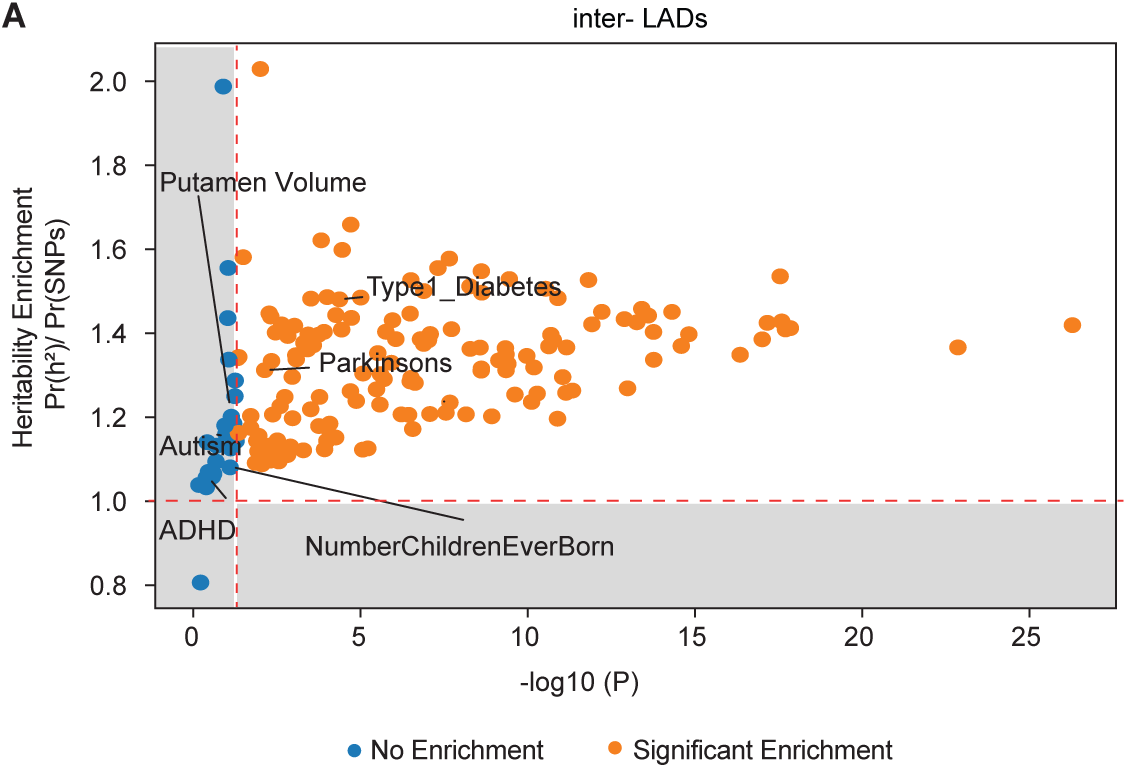
Genetic analysis inter-LADs with complex neuropsychiatric disorder-associated variants. (**A**) Scatter plots showing the enriched per-SNP heritability from partitioned LDSC analyses for SNPs associated with genes from inter-LADs. X-axis: P values in minus log10 scale; Y-axis: Enrichment of heritability for each SNP. Each dot represents a GWAS phenotype tested and is color-coded by their significance of enrichment.

**Supplementary Figure 6.**
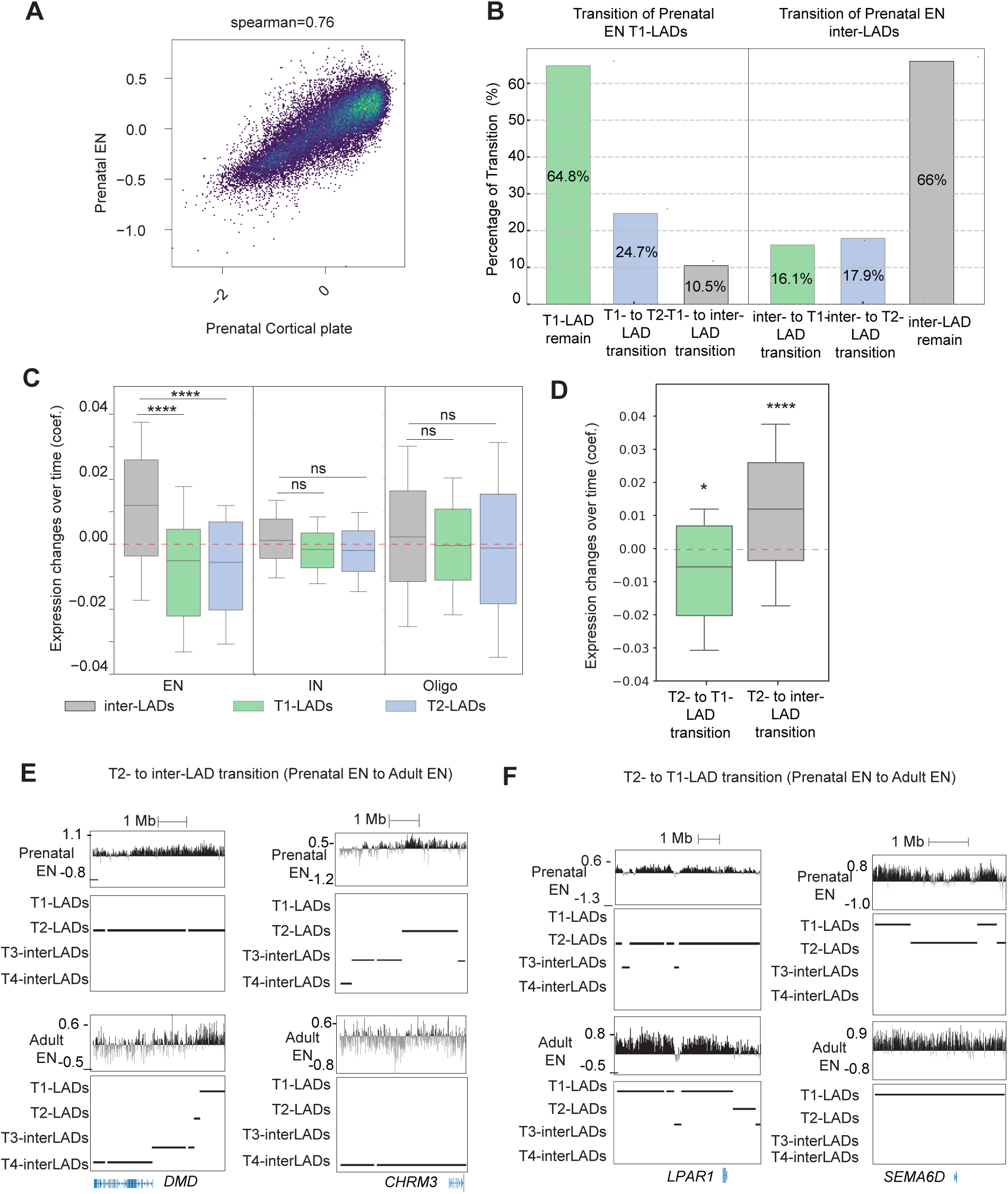
LAD substructure remodeling is accompanied by the relocation of genes involved in key biological functions during neuronal maturation. (**A**) Scatter plot comparing the LMNB1 enrichment across developing cortical plates and prenatal excitatory neurons (EN). (**B**) Bar plot showing the percentage of prenatal EN T1-LADs (left panel) or prenatal EN inter-LADs (right panel) that either remain stable or transition to T1-LAD or T2-LAD during neuronal maturation. Percentages of each transition state are indicated within the corresponding bars. (**C**) Boxplots showing genome-wide gene expression changes within excitatory neurons (ENs), inhibitory neurons (IN) and oligodendrocytes (Oligo) during neuronal maturation. Dashed red line represents the genome background. Statistical significance indicated as **** (p < 0.0001) or ns (not significant). (**D**) Boxplot showing gene expression changes over time for genes located in regions transitioning from T2-LADs to T1-LADs (green) or to inter-LADs (gray) during neuronal maturation. Dashed red line represents the genome background. Statistical significance: *p < 0.05; ****p < 0.0001. (**E**) Representative genome tracks highlighting transitions of gene locus from T2- to T1-LADs region during maturation of postmitotic excitatory neurons (left: chr9:106,569,266-112,394,032 region containing gene *LPAR1*, right: chr15:41,284,334-53,612,398 region containing gene *SEMA6D*. (**F**) Representative genome tracks highlighting transitions of gene locus from T2- to inter-LADs region during maturation of postmitotic excitatory neurons (left: chrX:31,050,024-35,803,159 region containing gene *DMD*, right: chr1:235,500,788-239,900,535 region containing gene *CHRM3.* Scale bar represents 1 Megabase (Mb).

## Methods

### Brain tissues samples

Five de-identified second-trimester human tissue samples were collected at the Zuckerberg San Francisco General Hospital (ZSFGH). Acquisition of second-trimester human tissue samples was approved by the UCSF Human Gamete, Embryo and Stem Cell Research Committee (study number 10-05113). All experiments were performed in accordance with protocol guidelines. Informed consent was obtained before sample collection and use for this study. Four adolescent tissue samples without known neurological disorders were obtained at the UCSF Pediatric Neuropathology Research Laboratory (PNRL) led by Dr. Eric Huang. These samples were acquired with patient consent in strict observance of the legal and institutional ethical regulations and in accordance with research protocols approved by the UCSF IRB committee (study number 13-11832). These samples were dissected and snap-frozen either on a cold plate placed on a slab of dry ice or in isopentane on dry ice.

### Tissues handling and generation of GO-CaT data

Microdissection was performed on prenatal human tissues to obtain the cortical plate region. Single cells were dissociated from cortical plates using the Papain Dissociation System (Worthington Biochemical). Cells were then fixed with methanol-free formaldehyde (Thermo Fisher Scientific) at a final concentration of 0.1% at room temperature for 2 minutes and quenched with 1.25 M glycine at room temperature for another 2 minutes.

Subsequently, cells were bound to BioMagPlus concanavalin A beads (Polysciences) and permeabilized using isotonic lysis buffer (20 mM Tris-HCl, pH 7.4, 150 mM NaCl, 3 mM MgCl2, 0.1% NP-40, 0.1% Tween-20, 1% BSA, and 1× protease inhibitors) on ice for 7 minutes. Cells were then incubated in antibody buffer (20 mM HEPES, pH 7.6, 150 mM NaCl, 2 mM EDTA, 0.5 mM spermidine, 1% BSA, and 1× protease inhibitors) containing either anti-LaminB1 antibody (Abcam, 16048, 1:100) or normal rabbit IgG antibody (Cell Signaling Technology, 2729, 1:100) at 4 °C overnight. The following day, cells were washed and incubated with antibody buffer containing guinea pig anti-rabbit secondary antibody (1:100, Novus Biologicals, NBP1-72763) at 4 °C for 1 hour. Cells were then washed twice using wash buffer (20 mM HEPES, pH 7.6, 150 mM NaCl, 0.5 mM spermidine, 0.01% digitonin, and 1× protease inhibitors) and incubated in high-salt wash buffer (20 mM HEPES, pH 7.6, 300 mM NaCl, 0.5 mM spermidine, 0.01% digitonin, and 1× protease inhibitors) containing pA-Tn5 (Epicypher) at 4 °C for 1 hour. Afterward, cells were washed twice with high-salt wash buffer and resuspended in the same buffer with the addition of 10 mM MgCl2. The resuspended cells were incubated in a PCR machine for 1 hour at 37 °C for tagmentation. After the reaction was complete, 1.7 μl of 0.5 M EDTA, 0.5 μl of 10% SDS, and 0.5 μl of 20 mg/mL proteinase K were added to the samples, followed by a 1-hour incubation at 55 °C. DNA was then extracted using phenol-chloroform. The extracted DNA was PCR-amplified using NEBNext High-Fidelity 2X PCR Master Mix (NEB) with a universal i5 primer and a unique i7 index. The amplified DNA was size-selected post-PCR using SPRIselect beads (Beckman Coulter) and eluted in TE buffer. GO-CaT library size and concentration were quantified using the High Sensitivity D1000 TapeStation assay (Agilent) and Qubit High Sensitivity dsDNA assay (Invitrogen). Finally, GO-CaT libraries were sequenced at the UCSF CAT Core using 150-bp paired-end sequencing on a NovaSeq X Plus system.

### Data processing of GO-CaT libraries

Sequencing reads were first trimmed to remove adapters and low-quality sequences using Trim Galore (https://www.bioinformatics.babraham.ac.uk/projects/trim_galore/) with default parameters. Then trimmed reads were mapped to the human reference genome (hg38) using Bowtie2 (v2.3.4.4). PCR duplicates were removed using Picard tools (http://broadinstitute.github.io/picard/). Properly paired reads with high mapping quality (MAPQ score > 30) were kept for further analysis. For visualization on the UCSC Genome Browser, the final bam files of LaminB1 were converted to bigwig files using bamCompare of deepTools (v3.4.1) with default parameters with IgG as the control bam files. For the visualization of histone modifications, bigwig files were generated using bamCoverage of deepTools (v3.4.1) with default parameters. For domain calling, the genome was binarized into 10kb bins and HMM domains were called using approach described previously^11^. Promoters were annotated using HOMER.

### Nuclei isolation, FANS and generation of EN GO-CaT data

Frozen cortex tissues were cut into small pieces and transferred to a pre-chilled 7-ml Wheaton™ Dounce Tissue Grinder containing 5 ml nuclei extraction buffer (NEB: 10 mM HEPES pH 7.4, 25 mM KCl, 5 mM MgCl2, 0.25 M sucrose, 0.1% Triton X-100, 1x Halt protease inhibitor cocktail (Thermofisher)). While still on ice, the tissue was dissociated with 5-6 strokes of loose pestle (A) followed by 8-10 strokes of tight pestle (B), until no tissue pieces were visible. The sample was incubated on ice for ∼5 min after which it was transferred to a pre-chilled 15-ml conical tube. The sample was centrifuged at 500g for 10 min at 4°C. The supernatant was removed, and the nuclear pellet was resuspended in 10-ml NEB without Triton X-100. The homogenate was passed through a 40μ-strainer into a 50-mL conical tube. Formaldehyde (Thermofisher) was added to a final concentration of 0.1% and the sample was incubated for 2 min at room temperature with rotation. Glycine was added to a final concentration of 75 mM to quench the reaction. BSA was added to a final concentration of 1% and the sample was centrifuged at 500g for 10 min at 4°C. The supernatant was discarded, and the nuclear pellet was resuspended in 1 mL staining buffer (PBS + 1% BSA). The nuclei were filtered through a 40-µ strainer into a 1.5-mL low-bind microcentrifuge tube and counted under the microscope. The nuclei were pelleted at 500g for 10 min at 4°C and resuspended in 100-150 μl of staining buffer.

Antibody staining was carried out for overnight at 4°C on rotation with the following antibodies: SATB2-Alexa Fluor 647 (Abcam, 1:100 dilution) and NeuN (ABN90, 1:300). The next day, nuclei were washed twice with PBS with 1% BSA, then stained with secondary antibody anti-Geuinea pig percp (Jackson ImmunoResearch, 1:100) for 1 hour at 4°C, 900 μl of staining buffer was added and the samples were centrifuged at 500g for 10 min at 4°C. The nuclear pellet was resuspended in 1 mL of staining buffer and centrifuged at 500g for 10 min at 4°C. Nuclei were resuspended in 1-2 mL of staining buffer (depending upon the starting material and yield) and filtered into a FACS (BD) tube. DAPI was added at 1μg/ml just before FANS. FANS was conducted on BD FACS Aria II Cytometer using a 70-μm nozzle. Sorted nuclei were collected in 5-ml tubes containing 300-500 μl of collection buffer (PBS + 5% BSA). Sorted nuclei were collected by centrifuging at 500g for 10 min at 4°C and incubated with Frab Fragment blocking the staining antibodies (Jackson Immuno) for 20mins on ice. The nuclei pellet was then washed twice with PBS containing 1% BSA and processed for GO-CaT.

### DNA-FISH and image analysis

DNA-FISH and image analysis was performed as described previously^14^. For the analysis of each condition of T1- or T2-LAD gene, 40–60 nuclei were analyzed.

### SNP Heritability Partitioning

We applied LDSC as previously described to partition SNP heritability for each neuropsychiatric trait using genes within prenatal T1-LAD, T2-LAD or inter-LAD regions^29^.

### Gene expression analyses

RNA-seq analysis of excitatory neurons were downloaded from public dataset^24^. Fastq files were aligned to the reference genome (GENCODE GRCh38) by STAR 2.7.11b after adaptor trimming. Reads that passed QC and with read length greater than 50 bp were carried over for downstream counting after mapping to a gene transfer format (GTF) file. Read counts were calculated using featureCounts. All differential expression analysis was performed by DESeq2 (v1.4) using raw counts across all conditions or cell types. Genes with read count > 50 in at least one sample of a comparison were kept for further analysis. A given gene was considered significantly changing if the false-discovery-rate (FDR) is < 0.05, P value is < 0.05, and Log1p fold-change is ≥ 0.05. Transcripts Per Million (TPM) were calculated to estimate the absolute expression level for each gene.

### Gene ontology analyses

Gene ontology (GO) analysis was performed using clusterProfiler (v3.19) with the default parameters. GO terms with FDR less than 0.05 were designated with significant functional enrichment.

### Statistics and data reproducibility

All the GO-CaT experiments in the human cortical plate or sorted neurons were performed for at least two biological replicates. No statistical test was used to predetermine the sample size used. P values were derived from Wilcoxon’s rank-sum test for comparisons throughout the study. Common domains of LADs/SPADs from two biological replicates were used for the analysis.

### Data and code availability

The datasets generated during the current study will be available on public repositories prior to the time of publication. The codes used in the data analyses will be available at GitHub.

## Acknowledgements

This work was supported by NIH grants R01NS112357, R01NS124881, R2NS1125978, P01 NS083513, VA grant I01BX000252, UCSF CIRM Scholars Training Program EDUC4-12812, the Chad Tough Foundation, and the Pathway for Breakthrough in Biomedical Research (PBBR), UCSF. The UCSF Parnassus Flow CoLab (RRID:SCR_018206) is supported in part by Grant NIH P30 DK063720 and by the NIH S10 Instrumentation Grant S10 1S10OD021822-01. Chujing Zhang was supported by UCSF-California Institute for Regenerative Medicine (CIRM) Scholars Training Program via education grant # EDUC4-12812. Li Wang was supported by NIMH grant K99MH131832. We thank Li Wang, Cheng Wang and members of Jingjing Li lab for providing suggestions on the experimental design and computational analysis for this study. We thank Evan R. Semenza for providing helpful suggestions for the manuscript. We thank flow cytometry, microscopy, and sequencing cores at UCSF.

## Author Contributions

Conceptualization, C.Z. and D.A.L.; Methodology, C.Z., L.W., M.L.; Formal Analysis, C.Z., M.L.; Investigation, C.Z. S.H.A, E.G, L.W., M.L.; Resources, E.J.H., J.L., A.R.K., and D.A.L.; Writing, C.Z., and D.A.L.; Supervision, J.L., A.R.K. and D.A.L.

## References

1. Belmont AS. Nuclear Compartments: An Incomplete Primer to Nuclear Compartments, Bodies, and Genome Organization Relative to Nuclear Architecture. Cold Spring Harb Perspect Biol 2022 Jul 1; 14(7).

2. Bickmore WA, van Steensel B. Genome architecture: domain organization of interphase chromosomes. Cell 2013 Mar 14; 152(6): 1270–1284.

3. Brueckner L, Zhao PA, van Schaik T, Leemans C, Sima J, Peric-Hupkes D, et al. Local rewiring of genome-nuclear lamina interactions by transcription. EMBO J 2020 Mar 16; 39(6): e103159.

4. Guerreiro I, Rang FJ, Kawamura YK, Kroon-Veenboer C, Korving J, Groenveld FC, et al. Antagonism between H3K27me3 and genome-lamina association drives atypical spatial genome organization in the totipotent embryo. Nat Genet 2024 Oct; 56(10): 2228–2237.

5. Peric-Hupkes D, Meuleman W, Pagie L, Bruggeman SW, Solovei I, Brugman W, et al. Molecular maps of the reorganization of genome-nuclear lamina interactions during differentiation. Mol Cell 2010 May 28; 38(4): 603–613.

6. Poleshko A, Shah PP, Gupta M, Babu A, Morley MP, Manderfield LJ, et al. Genome-Nuclear Lamina Interactions Regulate Cardiac Stem Cell Lineage Restriction. Cell 2017 Oct 19; 171(3): 573-587 e514.

7. Reddy KL, Zullo JM, Bertolino E, Singh H. Transcriptional repression mediated by repositioning of genes to the nuclear lamina. Nature 2008 Mar 13; 452(7184): 243–247.

8. Finlan LE, Sproul D, Thomson I, Boyle S, Kerr E, Perry P, et al. Recruitment to the nuclear periphery can alter expression of genes in human cells. PLoS Genet 2008 Mar 21; 4(3): e1000039.

9. Wang H, Xu X, Nguyen CM, Liu Y, Gao Y, Lin X, et al. CRISPR-Mediated Programmable 3D Genome Positioning and Nuclear Organization. Cell 2018 Nov 15; 175(5): 1405–1417 e1414.

10. van Steensel B, Belmont AS. Lamina-Associated Domains: Links with Chromosome Architecture, Heterochromatin, and Gene Repression. Cell 2017 May 18; 169(5): 780–791.

11. Shah PP, Keough KC, Gjoni K, Santini GT, Abdill RJ, Wickramasinghe NM, et al. An atlas of lamina-associated chromatin across twelve human cell types reveals an intermediate chromatin subtype. Genome Biol 2023 Jan 23; 24(1): 16.

12. Lui JH, Hansen DV, Kriegstein AR. Development and evolution of the human neocortex. Cell 2011 Jul 8; 146(1): 18–36.

13. Greil F, Moorman C, van Steensel B. DamID: mapping of in vivo protein-genome interactions using tethered DNA adenine methyltransferase. Methods Enzymol 2006; 410: 342–359.

14. van Schaik T, Vos M, Peric-Hupkes D, Hn Celie P, van Steensel B. Cell cycle dynamics of lamina-associated DNA. EMBO Rep 2020 Nov 5; 21(11): e50636.

15. van Steensel B, Henikoff S. Identification of in vivo DNA targets of chromatin proteins using tethered dam methyltransferase. Nat Biotechnol 2000 Apr; 18(4): 424–428.

16. Chen Y, Zhang Y, Wang Y, Zhang L, Brinkman EK, Adam SA, et al. Mapping 3D genome organization relative to nuclear compartments using TSA-Seq as a cytological ruler. J Cell Biol 2018 Nov 5; 217(11): 4025–4048.

17. Kumar P, Gholamalamdari O, Zhang Y, Zhang L, Vertii A, van Schaik T, et al. Nucleolus and centromere TSA-Seq reveals variable localization of heterochromatin in different cell types. bioRxiv 2023 Nov 1.

18. Ahanger SH, Delgado RN, Gil E, Cole MA, Zhao J, Hong SJ, et al. Distinct nuclear compartment-associated genome architecture in the developing mammalian brain. Nat Neurosci 2021 Sep; 24(9): 1235–1242.

19. Altemose N, Maslan A, Rios-Martinez C, Lai A, White JA, Streets A. muDamID: A Microfluidic Approach for Joint Imaging and Sequencing of Protein-DNA Interactions in Single Cells. Cell Syst 2020 Oct 21; 11(4): 354–366 e359.

20. Wang L, Wang C, Moriano JA, Chen S, Zuo G, Cebrian-Silla A, et al. Molecular and cellular dynamics of the developing human neocortex at single-cell resolution. bioRxiv 2024 Aug 4.

21. Rao SS, Huntley MH, Durand NC, Stamenova EK, Bochkov ID, Robinson JT, et al. A 3D map of the human genome at kilobase resolution reveals principles of chromatin looping. Cell 2014 Dec 18; 159(7): 1665–1680.

22. Lieberman-Aiden E, van Berkum NL, Williams L, Imakaev M, Ragoczy T, Telling A, et al. Comprehensive mapping of long-range interactions reveals folding principles of the human genome. Science 2009 Oct 9; 326(5950): 289–293.

23. Dekker J, Marti-Renom MA, Mirny LA. Exploring the three-dimensional organization of genomes: interpreting chromatin interaction data. Nat Rev Genet 2013 Jun; 14(6): 390–403.

24. Gibcus JH, Dekker J. The hierarchy of the 3D genome. Mol Cell 2013 Mar 7; 49(5): 773–782.

25. Heffel MG, Zhou J, Zhang Y, Lee DS, Hou K, Pastor-Alonso O, et al. Temporally distinct 3D multi-omic dynamics in the developing human brain. Nature 2024 Nov; 635(8038): 481–489.

26. Lee DS, Luo C, Zhou J, Chandran S, Rivkin A, Bartlett A, et al. Simultaneous profiling of 3D genome structure and DNA methylation in single human cells. Nat Methods 2019 Oct; 16(10): 999–1006.

27. Harris HL, Gu H, Olshansky M, Wang A, Farabella I, Eliaz Y, et al. Chromatin alternates between A and B compartments at kilobase scale for subgenic organization. Nat Commun 2023 Jun 6; 14(1): 3303.

28. Song M, Pebworth MP, Yang X, Abnousi A, Fan C, Wen J, et al. Cell-type-specific 3D epigenomes in the developing human cortex. Nature 2020 Nov; 587(7835): 644–649.

29. Herring CA, Simmons RK, Freytag S, Poppe D, Moffet JJD, Pflueger J, et al. Human prefrontal cortex gene regulatory dynamics from gestation to adulthood at single-cell resolution. Cell 2022 Nov 10; 185(23): 4428–4447 e4428.

30. Lange M, Bergen V, Klein M, Setty M, Reuter B, Bakhti M, et al. CellRank for directed single-cell fate mapping. Nat Methods 2022 Feb; 19(2): 159–170.

31. Buniello A, MacArthur JAL, Cerezo M, Harris LW, Hayhurst J, Malangone C, et al. The NHGRI-EBI GWAS Catalog of published genome-wide association studies, targeted arrays and summary statistics 2019. Nucleic Acids Res 2019 Jan 8; 47(D1): D1005–D1012.

32. Bulik-Sullivan BK, Loh PR, Finucane HK, Ripke S, Yang J, Schizophrenia Working Group of the Psychiatric Genomics C, et al. LD Score regression distinguishes confounding from polygenicity in genome-wide association studies. Nat Genet 2015 Mar; 47(3): 291–295.

33. Finucane HK, Bulik-Sullivan B, Gusev A, Trynka G, Reshef Y, Loh PR, et al. Partitioning heritability by functional annotation using genome-wide association summary statistics. Nat Genet 2015 Nov; 47(11): 1228–1235.

34. Dolphin AC, Lee A. Presynaptic calcium channels: specialized control of synaptic neurotransmitter release. Nat Rev Neurosci 2020 Apr; 21(4): 213–229.

35. Higgins D, Graupner M, Brunel N. Memory maintenance in synapses with calcium-based plasticity in the presence of background activity. PLoS Comput Biol 2014 Oct; 10(10): e1003834.

36. Kennedy MB. Synaptic Signaling in Learning and Memory. Cold Spring Harb Perspect Biol 2013 Dec 30; 8(2): a016824.

37. Wallace JL, Pollen AA. Human neuronal maturation comes of age: cellular mechanisms and species differences. Nat Rev Neurosci 2024 Jan; 25(1): 7–29.

38. Bron R, Vermeren M, Kokot N, Andrews W, Little GE, Mitchell KJ, et al. Boundary cap cells constrain spinal motor neuron somal migration at motor exit points by a semaphorin-plexin mechanism. Neural Dev 2007 Oct 30; 2: 21.

39. Nakanishi Y, Izumi M, Matsushita H, Koyama Y, Diez D, Takamatsu H, et al. Semaphorin 6D tunes amygdalar circuits for emotional, metabolic, and inflammatory outputs. Neuron 2024 Sep 4; 112(17): 2955–2972 e2959.

40. Yung YC, Stoddard NC, Mirendil H, Chun J. Lysophosphatidic Acid signaling in the nervous system. Neuron 2015 Feb 18; 85(4): 669–682.

